# DNA fingerprinting: an effective tool for taxonomic identification of processed precious corals

**DOI:** 10.1101/813865

**Authors:** Bertalan Lendvay, Laurent E. Cartier, Mario Gysi, Joana B. Meyer, Michael S. Krzemnicki, Adelgunde Kratzer, Nadja V. Morf

## Abstract

Precious coral species have been used to produce jewelry and ornaments since antiquity. Due to the high prices at which corals are traded, coral beds have been heavily fished. Hence, fishing and international trade regulations were put in place. However, poaching remains extensive and mislabeling of products is common. To this date, the control of precious coral exploitation and enforcement of trade rules have been largely impaired by the fact that species of processed coral skeletons can be extremely difficult to distinguish even for trained experts.

Here, we developed methods to use DNA recovered from worked precious coral skeletons to identify their species. We evaluated purity and quantity of DNA extracted using five different techniques. Then, a minimally invasive sampling protocol was tested, which allowed genetic analysis without compromising the value of the worked coral objects.

We found extraction of pure DNA possible in all cases using 100 mg skeletal material and over half of the cases when using “quasi non-destructive” sampling with sampled material amount as low as 2.3 mg. Sequence data of the recovered DNA gave a strong indication that the range of precious coral species present in the trade is broader than previously anticipated.

## Introduction

Precious corals are among the most appreciated and oldest known gems. They are valued for their color, purity and hard material, and have thus been collected and used for adornment for millennia ^1–3^. The strict coral harvesting and trading regulations introduced in recent decades along with growing demand in Asia, have led to an increase in prices of precious corals used in jewelry ^4–6^.

The precious coral material used for jewelry is the cut and polished hard coral skeletal axis, which is a biogenic material created by a biomineralization process^7^. In this process, closely packed magnesium-rich calcite crystals are secreted by coral polyps (1-2 mm in size) to build up a skeleton over decades. The polyps can thrive on the surface of the skeleton as colonies connected and surrounded by a 0.5-1 mm thick surface tissue (coenenchyme) ^8^. All deep-sea dwelling precious coral species belong to the family Coralliidae. The Coral Commission of The World Jewellery Confederation (CIBJO) lists eight Coralliidae species as significant in the precious coral jewelry industry ^9,10^. Precious coral products are sold worldwide with production centers located in Italy, Japan and Taiwan and large-scale trade of raw material between these areas ^5,6,11^.

Until recent decades, the populations of these highly coveted marine animals experienced exploitation in boom and bust cycles where the discovery of precious coral beds led to rushes by coral fishers and these beds were exploited as long as it remained economically feasible ^12,13^. The high price for precious corals made unreported fishing and poaching appealing for many ^14^. Therefore, local and international regulations were put in place to control both fishing and international trade of precious corals, among which four Pacific species were listed in Appendix III of the Convention on International Trade in Endangered Species of Wild Fauna and Flora (CITES) at the request of China ^4,13,15,16^ (Table 1). Despite these actions, illegal poaching and unregulated trafficking have remained frequent and extensive phenomena until today ^4,5,17^. It has also been reported that traders may often not be aware of the origin and species of their jewelry products or, moreover, deliberately mislabel their products ^4,6^. At the same time, consumers and jewelers increasingly request specific information about precious corals, particularly their geographic origin and species, mainly due to the perceptions of value that different types of coral have in the market and possible sustainability considerations ^18^.

**Table 1.** Distribution, CITES listing and trade names of the eight precious coral species considered relevant in the jewelry industry by The World Jewellery Confederation. Note that the different *Hemicorallium* species are sometimes suggested to be traded under the name *Pleurocorallium secundum* ^9,10,18,47^. Data compiled from ^6,9,10,18^

| Species | Distribution | CITES listed | Traditional trade names for sub-varieties of the species | Scientific listing in trade |
| --- | --- | --- | --- | --- |
| *Corallium japonicum*<br>Japanese red coral | West-Pacific | Yes | Oxblood, Red blood, Aka, Moro. | *Corallium japonicum* |
| *Corallium rubrum*<br>Mediterranean red coral | Mediterranean | No | Sardinian, Sardegna, Mediterranean, Sciacca | *Corallium rubrum* |
| *Hemicorallium laauense*<br>Deep-sea coral | North Pacific | No | Deep-sea Midway, Deep sea, New coral, Sensei | *Pleurocorallium secundum* |
| *Hemicorallium regale*<br>Garnet coral | North Pacific | No | Garnet | *Pleurocorallium secundum* |
| *Hemicorallium sulcatum*<br>Miss coral | West-Pacific | No | Miss, Missu, Misu | *Pleurocorallium secundum* |
| *Pleurocorallium elatius*<br>Pink coral | West-Pacific | Yes | Angel skin, Satsuma, Momo, Magai, Boké, Pelle d'angelo, Cerasuolo | *Pleurocorallium elatius* |
| *Pleurocorallium konojoi*<br>White coral | West-Pacific | Yes | Pure white, Shiro, Bianco | *Pleurocorallium konojoi* |
| *Pleurocorallium secundum* | North Pacific | Yes | Rosato, Midway, White/Pink | *Pleurocorallium secundum* |

Therefore, accurate taxonomic identification of precious coral products is of paramount importance for both efficient enforcement of precious coral trade regulations and for the jewelry industry. However, species of polished corals can be extremely difficult to distinguish even for trained experts based on morphological characteristics, and proper analytical tools to conclusively identify the species of worked precious corals are still lacking ^6,12,18,19^.

The various analytical methods tested to distinguish precious coral species based on skeletal material were either unable to provide clear-cut distinction among the different coral species (i.e. trace element analysis, ^20^; X-ray fluorescence spectroscopy and Raman spectroscopy, ^21^), or were not improved to become a standardized and easy-to-use tool i.e. immunolabeling, ^22^. As a novel approach, Cartier, et al. ^23^ recently proposed DNA analysis to distinguish species, assuming that coral DNA molecules can be trapped in the organic material or adhered to the CaCO_3_ crystals during the skeleton formation.

Genetic analyses have become a powerful analytical tool to elucidate the species identity and trace the geographic origin of various valuable artefacts of biogenic origin. These include processed products of tortoise shell ^24^, snake skin ^25^, fur ^26,27^, ivory ^28,29^ or tiger bones ^30^. Of greatest relevance to this present study, Meyer, et al. ^31^ reported quasi-nondestructive species identification of pearls based on DNA analysis, where so little amount of pearl material was used for the analyses that the gemological value of the pearl was not compromised. Particular biogenic materials require specific DNA extraction methods, moreover, we anticipate that DNA preserved in precious corals skeletons to be present in very small amounts and highly fragmented due to the lengthy skeleton-formation process and because the majority of corals are already dead when fished ^17,32–34^.

In the present study, we aim to explore whether precious coral skeleton fragments cut, carved and polished for jewelry could be taxonomically identified through genetic analysis. We compare five different DNA extraction methods to find the method producing the highest purity and quantity of DNA. We then apply the most successful technique to extract DNA using a minimally destructive sampling method and amplify and sequence the recovered DNA to taxonomically identify the coral samples. We demonstrate that genetic analysis of gem-quality precious corals is an efficient method to assess their species identity.

## Results

### Comparison of DNA retrieved from worked precious corals with five extraction methods

We evaluated which one of five candidate DNA extraction protocols is the most suited for precious coral skeletons. Each of the five tested methods (abbreviated as “W”, “F”, “B”, “E”, “Y”) have earlier proven to be useful to extract DNA from biomineralized material. DNA was extracted from each of a set of 25 worked coral skeletal samples with all five techniques, and DNA purity and quantity were assessed using real-time quantitative PCR (qPCR) technology.

To test DNA extract purity, we assessed PCR inhibition with qPCR using an internal amplification control molecule. Three extraction methods, “F”, “E” and “Y”, resulted in DNA with no detectable PCR inhibition effect from any of the tested 25 samples (Fig. 1, Supplementary Results S1). In contrast, a PCR inhibition effect was observed in 15 out of 25 samples extracted with the “B” method. Of these, complete inhibition of the PCR was observed in one case. Inhibition was also detected in three DNA extracts produced with the “W” method. Of these, no PCR product was observed at all in one sample.

**Figure 1.**
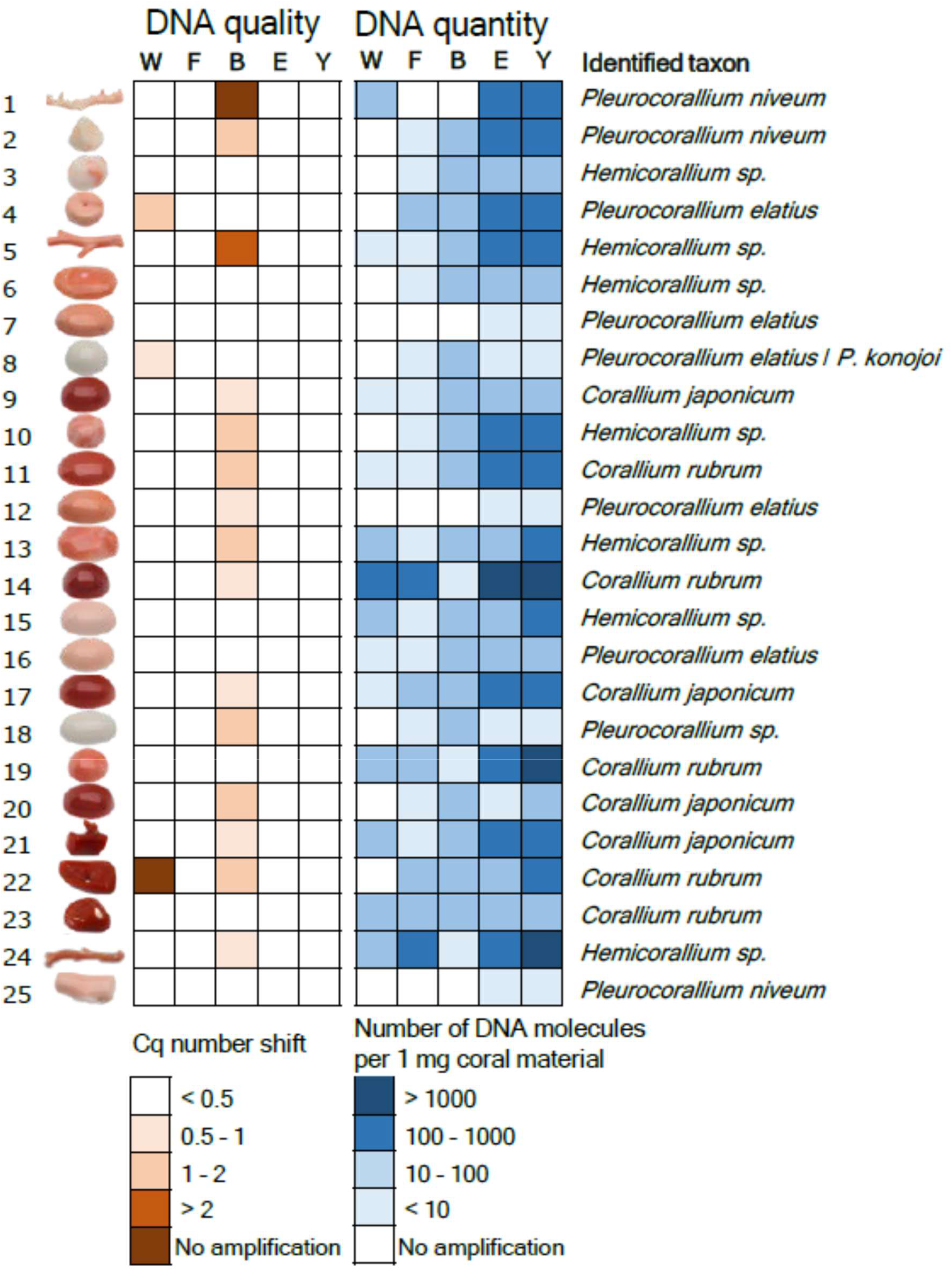
Results of the DNA extract purity and quantity measurement experiment and taxonomic identification of 25 worked precious coral samples. Five methods were used to extract DNA from equal amounts of material from each sample. PCR inhibition measurement and absolute template quantification was performed with quantitative real-time PCR. Two short mitochondrial DNA fragments were sequenced and each specimen was taxonomically assigned.

Absolute quantity of the DNA obtained with the five extraction techniques was tested using qPCR with a standard curve from a dilution series of a standard template DNA molecule with known concentrations. Throughout these analyses, the average qPCR efficiency was 88.5 % (± 3.6% standard deviation) and coefficient of determination for the calibration curve was R^2^ = 0.9947 (± 0.0035 standard deviation), respectively.

The five extraction methods yielded highly varying amounts of DNA (Fig. 1, Supplementry Results S1.). Methods “E” and “Y” both yielded PCR amplifications for all 25 samples. Method “W” yielded PCR product for 13 samples, while methods “F” and “B” both yielded PCR product for 21 samples. Overall, there was concordance among the amplification results; the 13 samples that amplified with method “W” also amplified with methods “F” and “B”, and the latter two methods amplified DNA of the very same 21 samples. Strong significant correlation was found between the copy numbers obtained from the same coral items with the “E” and “Y” methods (r=0.97, t=19.223, df=23, p<0.001). The DNA yield was higher with method “Y” than with method “E” (595 versus 944 molecules per mg coral skeleton with “E” and “Y”, respectively; paired t-test: t = - 2.8832, df = 24, p = 0.008). Focusing on the best performing “Y” method, DNA concentrations ranged three orders of magnitude; three samples had over 10^3^ DNA copies in each mg of skeleton material. In contrast, in five other samples this value was below 10 (Fig. 1).

### DNA extraction with “quasi non-destructive” sampling of worked precious coral skeletons

We developed a “quasi non-destructive” technique to take material for analysis from the worked corals with minimal weight loss and virtually invisible effects of the sampling (Fig. 2). A new set of 25 worked coral samples were sampled in this manner; removed material amounts ranged from 2.3 mg to 13.1 mg and were 7.9 mg on average. Modifications were applied to the lysis step of the “Y” extraction method compared to the original protocol, which resulted in an essentially complete dissolution of the coral powder. This allowed the amount of DNA that remained trapped in the undissolved powder to be kept to the minimum. Out of the 25 “quasi non-destructively” sampled worked coral objects, 16 gave qPCR amplicons at least twice (Fig. 3, Supplementary Results S1). Another two samples produced amplification only once and were omitted from further analyses. DNA copy numbers calculated per mg of coral skeleton were in the same range as in the case of the extractions carried out from c. 100 mg material using the “Y” method. However, the presence of unsuccessful amplifications and lower average copy number (160 DNA copies) recovered per mg of coral skeleton indicates that DNA recovery from low amount samples is less effective than from standard material amount, despite the amendments made in the DNA extraction protocol.

**Figure 2.**
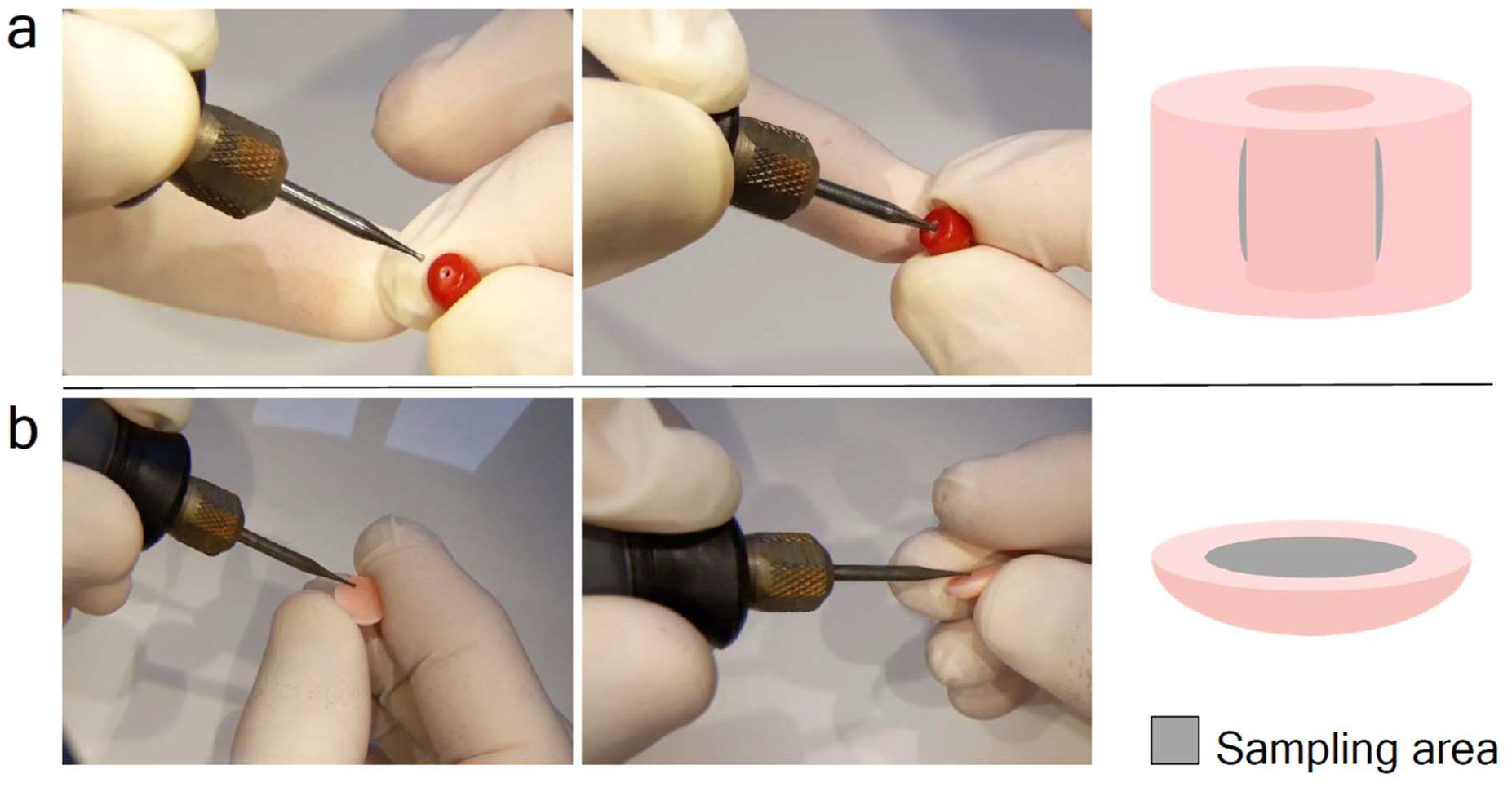
“Quasi non-destructive” sampling of worked coral skeletons. a) Widening the inner surface of existing drill-holes. b) Sampling the back side of items without an existing drill-hole.

**Figure 3.**
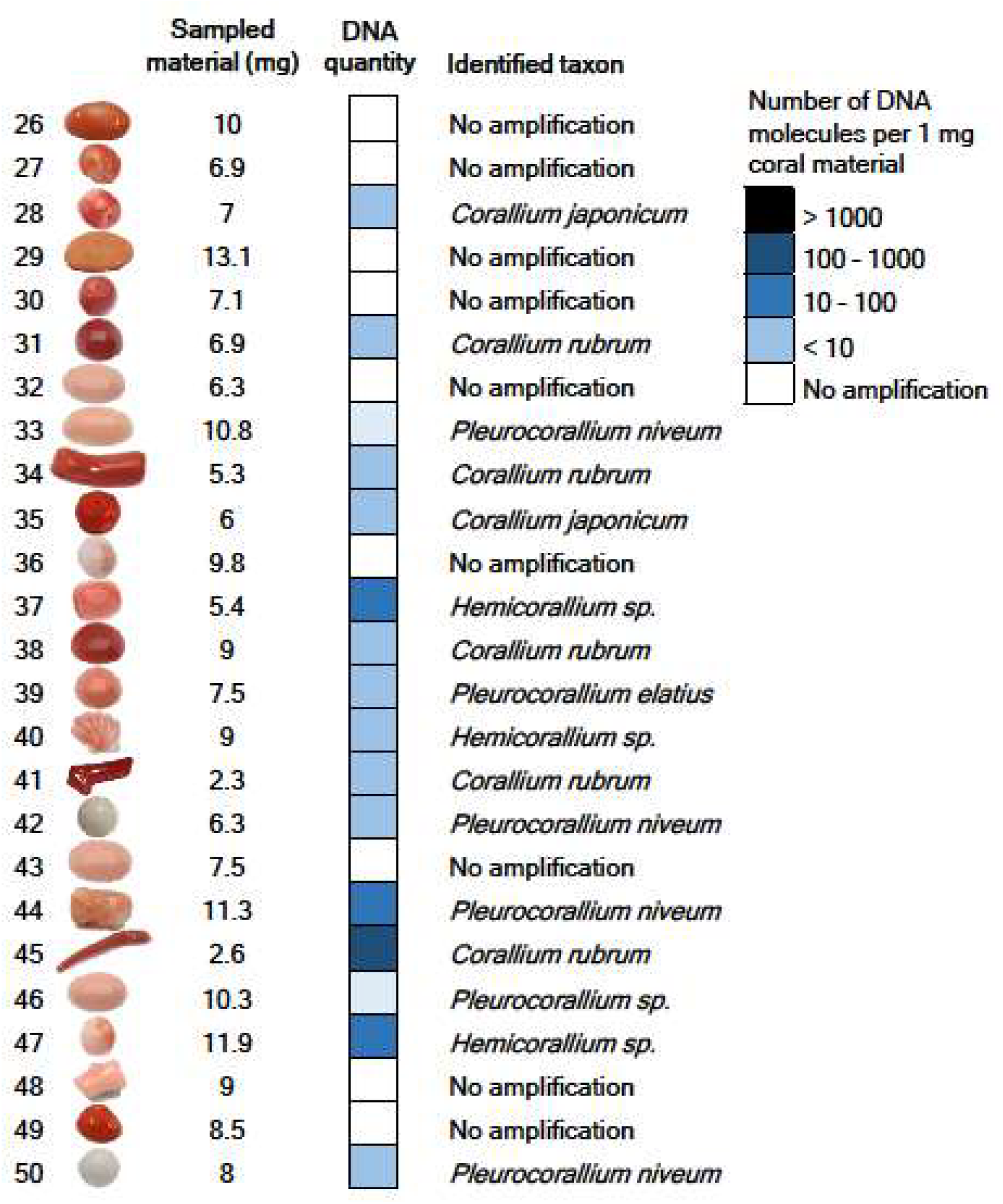
Results of DNA quantity measurement and taxonomic identification of 25 worked precious corals sampled by the minimally invasive technique. Absolute template quantification was performed with quantitative real-time PCR. Two short mitochondrial DNA fragments were sequenced and each specimen was taxonomically assigned.

### Taxonomic assignment of worked precious corals

We sequenced amplicons of the large ribosomal RNA gene subunit (LR) and the putative mismatch repair protein (MSH) fragments originating from a total of 41 worked coral skeletons. In our entire DNA sequence dataset, sequence of three OTUs did not align with neither with the LR nor the MSH reference sequences. NCBI BLAST search did not find any sequence entries in the NCBI database with higher than 95% sequence similarity to any of these sequences.

Length of concatenated LR and MSH sequences were between 264 base-pairs (bp) and 290 bp long per coral sample (Supplementary Results S2). Phylogenetic analysis identified 10 samples (11, 14, 19, 22, 23, 31, 34, 38, 41, 45) as *Corallium rubrum*, of which nine had sequences identical to either of two the reference *C. rubrum* sequences, and one (11) had a single variable site (Fig. 4). Six samples (9, 17, 20, 21, 28, 35) were identical with reference samples of *Corallium japonicum*, but also with the reference samples of *C. nix* and *C. tortuosum*.

**Figure 4.**
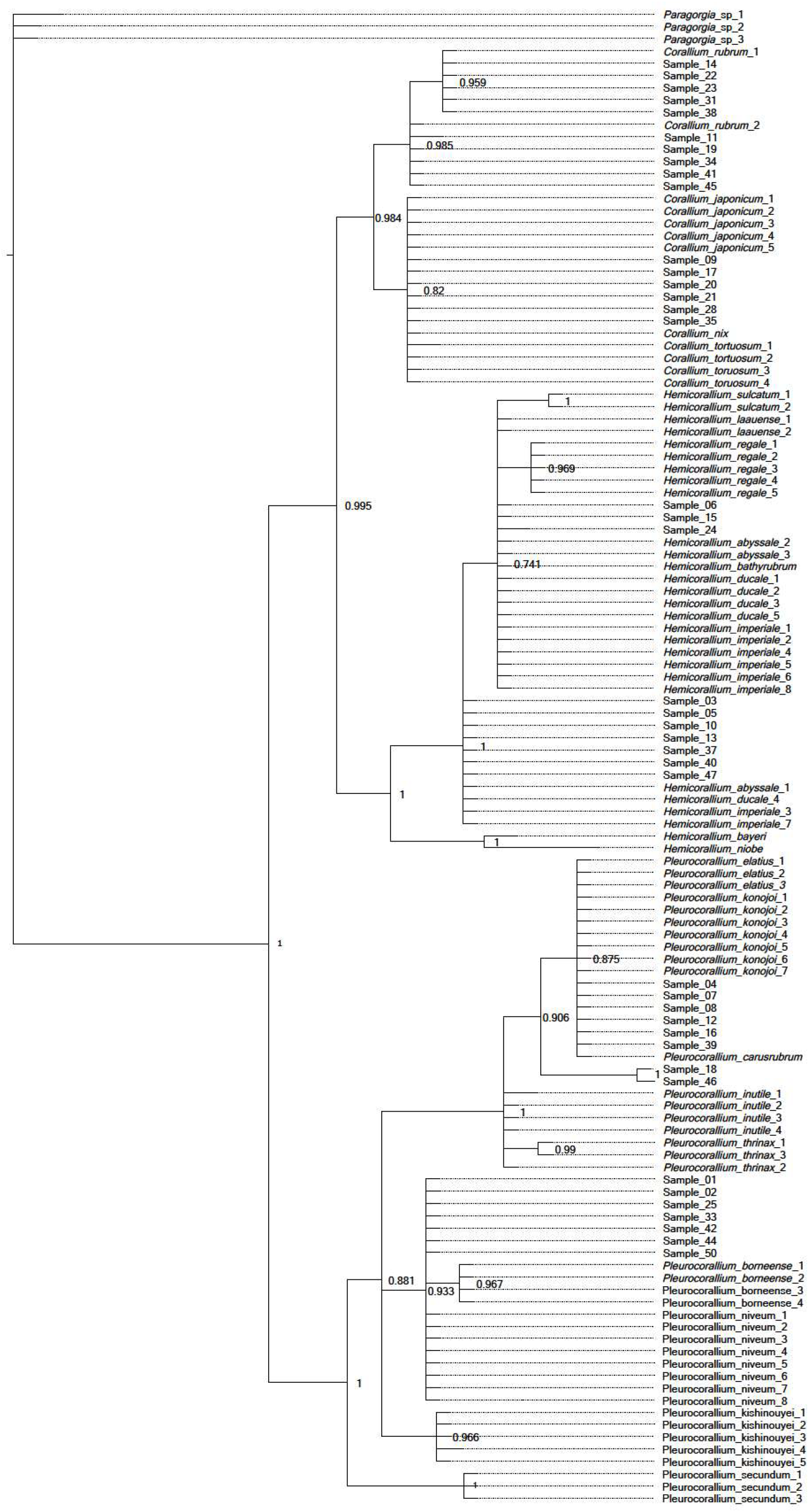
Majority-rule Bayesian phylogenetic tree constructed from combined mitochondrial LR and MSH region DNA sequence data of worked precious corals and reference samples. Posterior probability value is displayed after each tree node.

Three samples (6, 15, 24) formed a polytomic clade with Hemicorallium reference sequences. Two of these (6, 15) had sequences identical to *Hemicorallium laauense*, but also to samples of *H. abyssale*, *H. bathyrubrum*, *H. ducale* and *H. imperiale*. The third sample (24) was one bp different from these sequences. Seven samples (3, 5, 10, 13, 37, 40, 47) with identical sequences appeared as an unresolved clade basal to the formerly mentioned samples. These had identical sequences with *H. abyssale*, *H. ducale* and *H. imperiale*.

Six samples (4, 7, 8, 12, 16, 39) had identical sequences with *Pleurocorallium carusrubrum*, *P. elatius* and *P. konojoi* reference samples. Two samples (18, 46) formed a sister clade to the former group with the posterior probability value 1. Finally, seven identical samples (1, 2, 25, 33, 42, 44, 50) were same as sequences of *Pleurocorallium niveum*. These were grouped together as an unresolved tree branch.

## Discussion

Technical advancements and the growing body of reference DNA data have made genetic analyses a powerful tool to combat poaching, illegal trading and mislabeling of animal products ^35^. Application of genetic barcoding was suggested by Ledoux, et al. ^36^ as a forensic tool to identify species of corals. Acknowledging that the discriminatory power of standard species barcoding markers (e.g. the cytochrome c oxidase subunit I gene) is poor to distinguish the closely related precious coral species, these authors suggested development of custom designed species identification markers. Moreover, if the aim is to distinguish coral skeleton samples, then the high portion of fragmented DNA will call these markers to be as short as possible. A further challenge is if sampling of the coral skeleton is to be done with minimal material loss, and as consequence, the chosen DNA extraction method has to be capable of recovering DNA from small sample amount.

In our quest to find an optimal method to recover DNA from worked coral skeletons, we tested the performance of five DNA extraction methods, each on equal amount of coral material from the same set of 25 worked coral samples. We found two methods, protocol “E” and “Y” that yielded DNA that was successfully amplified and sequenced from all of the 25 tested corals. Methods “E” and “Y” are two similar techniques developed for the extraction of DNA from ancient eggshells and ancient bones. They only slightly differ in their lysis buffer ingredients and the type of DNA-binding silica column used for the purification of the recovered DNA molecules ^37,38^. These methods produced similar amounts of DNA, however method “Y” produced slightly higher DNA yield, particularly in the samples that had < 50 DNA copies per mg coral powder. The three other tested DNA extraction methods did not result in amplifiable DNA from all samples, which may be due to their inability to recover DNA coupled with PCR-inhibitory effect of co-extracted substances, which was detected in some extracts, PCR inhibition was not detected in any extracts produced with methods “W”, “E” and “Y”. By using these methods, PCR inhibition seems to be overcome in precious corals, unlike in other types of corals, where it led to technical challenges ^39^.

DNA concentration of the extracts differed largely; while in certain samples <10 copies per mg material was recovered, in some others this reached up to the order of magnitude of 10^3^ copies per mg material. The large variation in DNA preservation of the samples may be determined by their varying ages: corals are often fished decades after their death ^17,32,33^ and coral skeletons maybe stored for long before they get processed ^6^. However, without specific knowledge about the age of the samples this remains hypothetical.

Our test to choose the best DNA extraction protocol from potential methods was based on 100 mg of coral skeleton, which is a standard amount used for extracting DNA from pulverized material with the applied protocols. The essence of precious material testing would be to use as little material as possible, ideally using a “quasi non-destructive” sampling method. This means that the sampling area is not visible and the sampling does not cause significant weight loss of the coral object. Worked coral skeletons can be separated into two main types; the ones that have a hole drilled through the item (generally those that are strung for bracelets or necklaces) and the ones that do not have a hole, instead generally have flat reverse or bottom sides (those that are mounted to a frame and used as pendants, i.e. cabochons, or the carved figures used as ornaments). We performed “quasi non-destructive” sampling using a drill with a 0.8 mm diameter diamond engraver head taking care not to heat up the sampled object (no hard pressing of the drill and regular pauses to let the drill head cool down). With careful handling, it was possible to take sample material by slightly widening the internal surface of the ca. 1 mm wide drill-holes completely invisible by eye. From the cabochons, a thin layer was removed from the reverse side; therefore the visible front side remains unaffected by the sampling. Assuming approximately 3.8 kg/dm^3^ density of the precious corals ^9^, the removed 2.3-13.1 (in average 7.9 mg) mg sampled powder per sample corresponds to a 0.7 – 3.5 (2.1) mm^3^ volume loss of the items.

We were able to repeatedly produce PCR products for altogether 16 out of the 25 “quasi non-destructively” sampled worked coral skeletons. We cannot determine a threshold for the minimum amount of material necessary for successful genetic testing; the two samples processed with the lowest weight of coral powder, 2.3 mg and 2.6 mg, respectively, both produced results. Although it was not possible to genetically analyze all samples with the minimally destructive method, there might be a good chance that when analyzing several samples from a batch of samples, at least some will produce results.

We expected that the DNA sequences we generate will cluster together with reference sequences of one of the eight species listed by CIBJO as relevant in the jewelry industry. Our taxonomic identification markers allowed us to distinguish all of these eight precious coral species from the others with the exception of *Pleurocorallium elatius* and *P. konojoi*. However, unexpectedly, we found a much higher diversity within our samples, with several of our sequences not grouping together with the reference sequences of the eight species. Hence, we repeated the phylogenetic analysis with an extended reference sample set. The results of this analysis show that samples could clearly be identified as *Corallium rubrum*. The samples grouping together with *Corallium japonicum* also grouped together with two other species, *C. nix* and *C. tortuosum*, which, however, have white and pink color, respectively, unlike the dark red color of *C. japonicum* ^44,40^. Hence, we confidently identify these red corals as *C. japonicum*.

Samples that grouped together with the *Hemicorallium* references all had identical sequences with multiple *Hemicorallium* species. As a consequence, these samples could be identified only to the genus level as *Hemicorallium*. A part of these samples (i.e. 3, 5, 10, 13, 37, 40, 47) did not cluster with the three reportedly fished *Hemicorallium* species (*H. laauense*, *H regale*, *H. sulcatum*), but instead had identical sequences to other species (*H. abyssale*, *H. ducale* and *H. imperiale*) that all occur around the Hawaii islands, a historically important fishing area ^44,45,47^. This result strongly suggests that *H. laauense*, *H regale* and *H. sulcatum* are not the only *Hemicorallium* species appearing on the jewelry market.

Some samples had identical sequences of the three, both genetically and morphologically, very similar species, *Pleurocorallium. carusrubrum*, *P. elatius* and *P. konojoi* ^41,47^. Of these species, the latter two are well known in the jewelry industry, while the former is a recently described species known from single area of the West Pacific ^41^. To distinguish these species, the coloration of the skeletal axis may give a partial solution. In particular, the color of *P. carusrubrum* is red, *P. elatius* varies from white to dark pink, while *P. konojoi* is always pure white ^4,6,9^. Consequently, our specimens identified as one of these species with pink shading may be identified as *P. elatius*, while our samples with white color are determined as *P. elatius*/*P. konojoi*.

Of our multiple samples within the Pleurocorallium clade that did not group together with the species traditionally accepted as present on the coral market (*P. elatius*, *P. konojoi* and *P. secundum*), two samples (18, 46) formed an individual clade and were identified to the genus level as *Pleurocorallium*. DNA sequences of the other samples were all identical with the sequences of the *Pleurocorallium niveum* samples. This species was described from waters surrounding the Hawaii islands, which is a historically important coral fishing area ^42,43^.

The 41 samples that we managed to genetically analyze from 50 samples of a single collection is not representative to draw conclusions about the entire jewelry industry, but it indicates that there may be more species present in the trade than the eight precious coral species commonly listed as part of the jewelry industry (cf. ^9,10,18^). This is conceivable, if we consider that in the Pacific Ocean different precious coral species may co-occur and coral fishing does not seek to individually separate them based on species.

## Conclusions

This study is a proof of concept that genetic analysis can be an effective tool to taxonomically identify precious corals worked for jewelry. We demonstrated that while 100 mg coral skeletal material is sufficient for successful DNA extraction in all cases, DNA sequencing and taxonomic assignment were possible with minute amounts of “quasi non-destructive” samples in more than half of the cases. Among the worked precious corals examined in this study, DNA sequence analyses revealed several samples belonging to precious coral species previously not considered to be present in the jewelry industry. Future research should focus on broadening the reference data by sequencing multiple specimens for each species identified by experts to substantiate their intra- and interspecific genetic diversity. This will be an essential step in developing genetic tests to become a reliable and standardized method to promote sustainable use of precious corals in the jewelry industry.

## Materials and Methods

### Study species

The precious corals relevant in the high-end jewelry industry are Octocorallid Anthozoans that belong to the Alcyonacea order and Coralliidae family. Recent phylogenetic studies confirmed the existence of three genera in the family; *Corallium*, *Hemicorallium* and *Pleurocorallium* ^44,45^. Of the eight species listed by CIBJO as significant in the precious coral industry, a single, *Corallium rubrum*, is distributed in the Mediterranean Sea and has been fished since antiquity ^6^. Four other species, *Corallium japonicum*, *Hemicorallium sulcatum*, *Pleurocorallium elatius* and *Pleurocorallium konojoi* have been fished in the Western Pacific ocean since the early 19^th^ century ^12^. The remaining three species, *Hemicorallium laauense*, *Hemicorallium regale* and *Pleurocorallium secundum* were discovered on seamounts surrounding the Hawaii archipelago and were fished in large quantities during the second half of the 20^th^ century ^46^. Distribution, CITES listing and trade names of the eight precious coral species relevant in the jewelry industry are summarized in Table 1, while further details on their distribution, taxonomy, harvesting and conservation are presented in Supplementary Table S3.

### Genetic markers used in the study

We expected that the DNA extracted from the coral skeletons would be highly degraded. Therefore, we used markers developed on the mitochondrial genome, which is present in each cell in multiple copies and offers the best chances of achieving positive results for fragmented DNA. Octocoral mitochondrial genomes have an exceptionally low rate of evolution and standard taxonomic markers are unable to distinguish closely related species ^48,49^. Hence, we developed two genetic markers with the criteria that they are, at the same time, short to be suitable for degraded DNA and highly variable to maximize our ability to identify the precious coral species to the lowest possible taxonomic level. We expected each analyzed sample to originate from one of the eight precious coral species listed by CIBJO, thus developed our markers with the aim that they are capable of distinguishing these eight species. The two markers were used in combination to taxonomically identify the coral skeleton samples. Furthermore, one of them was used to test purity and DNA quantity of our extracts.

The two mitochondrial markers were developed based on DNA sequence data of Tu, et al. ^45^, which is the most detailed study on precious coral phylogeny to this date. Marker selection and procedure of designing PCR primers are detailed in Supplementary Methods S4.

Following examination of the phylogenetic resolution of multiple short mitochondrial genome fragments, we developed the two set of primers for the large ribosomal RNA gene subunit (LR) and the putative mismatch repair protein (MSH), respectively. The LR marker was used for the assessment of DNA extract purity and DNA quantification. Phylogenetic analysis using the LR and MSH markers showed that these two short markers were able to reconstruct the phylogenetic relationships obtained by much longer sequences, and they allowed the distinction of each of the eight precious coral species from each other, except for *Pleurocorallium elatius* and *P. konojoi*, which are not possible to conclusively distinguish based on the data of Tu, et al. ^45^ (Supplementary Methods S4).

### Comparison of DNA purity and quantity extracted with different methods

#### DNA extraction

All laboratory work was carried out at the Forensic Genetics department of the Institute of Forensic Medicine, University of Zurich, in the laboratory facility dedicated to human and animal forensic casework. We strictly adhered to the ISO 17025 guidelines throughout the laboratory workflow with stringent rules to avoid contamination and authenticate our results (Supplementary Methods S5). Precious coral samples used in this study originated from the collection of the Swiss Gemmological Institute SSEF, Basel, Switzerland.

Twenty-five worked coral samples were selected for the experiment (named samples 1-25, Supplementary Table S6). The samples were cleaned as described in Supplementary Methods S5 and crushed in a metal mortar with a metal pistil to produce crude coral powder, which was then transferred to a porcelain mortar and ground to fine powder. The coral skeleton powder was divided into five aliquots of equal weight, 100 mg ± 1 mg in general, except for four samples that had less available powder (Supplementary Table S6). The powder aliquots were used to extract DNA using five different extraction methods, which have proven to successfully recover DNA from biomineralized material (Table 2). For each method, we followed the protocols cited in Table 2. All DNA extracts were eluted in 100 µl and stored at - 20 °C.

**Table 2.** The five different methods tested to extract DNA from precious coral skeletons worked for jewelry. The abbreviated method name is followed by references of relevant studies where the method was used to extract DNA from calcified material, the chemical composition of the demineralization buffer, the method used for DNA binding and DNA purification and, finally, the exact protocol followed.

| Method | Previously used for | Demineralization | DNA binding | DNA purification | Protocol followed |
| --- | --- | --- | --- | --- | --- |
| “W” method | ancient bones <sup>50</sup>,<br>snail shells from museum collection <sup>51</sup> | EDTA solution with Proteinase-K | On celite particles in the presence of GuSCN containing buffer | Wizard PCR Preps DNA Purification System (Promega, Madison, WI, USA) | “WSU fast” method by Villanea, et al. <sup>51</sup> |
| “F” method | marine pearls <sup>31</sup> | EDTA solution | On spin column of the FastDNA SPIN Kit for Soil (MP Biomedicals, Illkirch-Graffenstaden, France) |  | Meyer, et al. <sup>31</sup> |
| “B” method | bones and teeth | PrepFiler BTA Lysis Buffer (Thermo Fisher, Waltham, MA, USA) with DTT and Proteinase-K | On magnetic particles of the PrepFiler BTA Forensic DNA Extraction Kit (Thermo Fisher) |  | vendor’s protocol |
| “E” method | subfossil eggshells <sup>38,52,53</sup> | Tris-EDTA solution with Triton X-100, DTT and Proteinase-K | On the spin column of the QIAquick PCR Purification Kit (Qiagen, Hilden, Germany) |  | Oskam and Bunce <sup>38</sup> |
| “Y” method | subfossil bones <sup>37</sup>,<br>subfossil and modern clam shells <sup>54</sup> | EDTA solution with N-laurylsarcosyl and Proteinase-K | On the spin column of the MinElute PCR Purification Kit (Qiagen) |  | “Y” method as described by Gamba, et al. <sup>37</sup> |

Abbreviation of chemical names: EDTA: Ethylenediaminetetraacetic acid, GuSCN: Guanidinium thiocyanate, DTT: Dithiotreitol

#### Assessment of the purity of the DNA extracts

We used qPCR to compare the purity of the DNA extracts produced from precious coral skeletons with five different extraction protocols. DNA purity was measured by testing the PCR inhibiting effect of the coral extracts during amplification of an internal positive control DNA fragment. We used 10^3^ copies of a synthetic oligonucleotide (gBlocks Gene Fragments; International DNA Technologies, Coralville, IA, USA ^55^) as internal amplification control (IAC, Supplementary Methods S7). The 197 bp sequence of the IAC matched 151 bp of the *C. rubrum* LR gene fragment (with manual introduction of five unique mismatches for contamination detection purpose) flanked by potato-specific sequences as primer sites following Nolan, et al. ^56^.

Following optimization (see Supplementary Methods S7), reactions were conducted in 20 µl volumes containing 1 × PowerUp SYBR Green Master Mix (Thermo Fisher), 1 µl of both 15 uM concentration primers, 10^3^ copies of the AIC in 3 µl and 3 µl coral DNA extract. Alongside the samples containing coral DNA extracts, we run three positive standard reactions that did not contain coral DNA. Following the manufacturer’s recommendation, reactions commenced with 50 °C for 2 minutes, which was followed by initial denaturation at 95 °C for 2 minutes and 50 cycles of denaturation at 95 °C for 15 seconds, primer annealing at 60 °C for 15 seconds and elongation at 72 °C for 1 minute. A melting-curve analysis was performed at the end of the reaction by heating the PCR products from 60 °C to 95 °C with 1% ramping speed. Each coral extract was run in triplicates on an ABI 7500 qPCR instrument (Thermo Fisher).

The quantification cycle (Cq) value of each reaction containing coral DNA extract was compared to the average Cq value of the tree positive standard reactions and then the three Cq shift values of each sample were averaged. Intensity of PCR inhibition in each reaction was determined as follows: we considered inhibition to be present if there was a 0.5< cycle Cq shift compared to the positive standard Cq. Four categories of PCR inhibition were considered: 0.5-1, 1-2, 2< cycle shifts and complete inhibition in case at least one out of the three reactions produced no PCR product.

#### Absolute DNA quantification of the coral DNA

Absolute quantification of the coral LR gene fragment was conducted by qPCR of the coral DNA using a calibration curve prepared as a series of standard reactions with known template DNA amount. The standards contained seven different 10-fold diluted template inputs (10^7^-10^1^ copies) of a GBlocks synthetic oligonucleotides of the 154 bp long sequence of the LR gene fragment characteristic to *C. rubrum* (with manual introduction of three unique mismatches for the purpose of contamination detection) flanked by the LR primer sequences (Supplementary Methods S7). Following optimization of the reaction setup (Supporting Methods S7), reactions were carried out in 20 µl volumes containing 1 × PowerUp SYBR Green Master Mix (Thermo Fisher), 1 µl of both 15 µM concentration primers and 3 µl coral DNA extract. The cycling conditions were identical as for the DNA extract purity test, except for adjusting the annealing temperature to 56 °C.

PCR was considered successful if at least two reactions of the triplicates amplified. Ct values were averaged for each sample and template molecule amount values were recalculated to number DNA molecules per mg of coral skeletons for each sample based on the volume of template, the DNA extract elution volume and the DNA extraction starting material amounts. We compared the DNA quantities gained with the extraction methods for which DNA was successfully amplified for all 25 samples with a correlation test and paired t-test in R ^57^.

### “Quasi non-destructive” sampling, DNA extraction and quantification

In the previous experiment, 25 samples were completely pulverized and five DNA extractions were carried out with different methods from each. The aim was to select the most suitable technique for extracting DNA from coral skeletons. In the current experiment, the best performing DNA extraction technique was used with “quasi non-destructive” sampling of processed corals. We define “quasi non-destructive” sampling as taking material for analysis from the worked objects without compromising its gemological value. A new set of 25 worked coral samples were selected from the SSEF coral collection for this experiment (named samples 26-50, Supplementary Table S8), and each was thoroughly cleaned as described in Supporting Methods S3. Two main types of samples were sampled differently: *i)* beads with drill-holes: the inner surface of the drill-hole was carefully widened (Fig. 2a); *ii)* worked items with no existing drill-hole: a small layer of the surface of the back side was removed (Fig. 2b). We used 0.8 mm diameter diamond engraver bit heads attached to a Dremel 4000-4 rotary tool (Dremel, Racine, WI, USA). Rotation speed was set to 10,000 rpm and coral drill-powder was left to drop in 1.5 ml collection tubes.

DNA was extracted from the quasi non-destructively sampled drill-powder of the 25 samples with the “Y” method. The material amount obtained by the “quasi non-destructive” sampling was far lower than the 100 mg used in the experiment comparing extraction methods, therefore we slightly modified the “Y” protocol to accommodate it to the low material amount. In particular, 200 µl lysis buffer was added to the coral powder and incubated at 56 °C for one hour with mixing, then another 100 µl lysis buffer was added. The lysis-mixture was then incubated again with mixing at 56 °C for one hour and then at 37 °C for additional 65 hours. The lysate was then mixed with 450 µl 1 × TE buffer and 3750 µl PB buffer (Qiagen) and the entire volume of the mixture was centrifuged through a MinElute (Qiagen) column, which washed then washed with PE buffer and the DNA was eluted in 35 µl EB buffer (Qiagen).

### Taxonomic identification

#### DNA amplification and sequencing

We sequenced the qPCR products of the LR fragment generated for the DNA quantity assessment. From each sample, one of the triplicates was selected for sequencing. The MSH region was then amplified for all 25 DNA samples extracted with the “Y” method in our DNA extraction test and the 16 DNA extracts from the “quasi non-destructive” sampling that gave amplification products for the LR region. The MSH was amplified in singlicate for each sample with identical reaction setup and cycling conditions as described above for the LR region.

The 16S and MSH PCR products were purified with the AMPure bead system (Beckman Coulter, Brea, CA, USA) and quantified with a Qubit 4 Fluorimeter (Thermo Fisher). The two amplicons of each DNA sample were pooled with equimolar concentrations, and sequencing libraries were constructed with the Ion Plus Fragment Library Kit (Thermo Fisher) according to the vendor`s protocol. The libraries were quantified with the Ion Library TaqMan Quantitation Kit (Thermo Fisher) and all samples were pooled with equimolar concentrations. Sequencing was carried out on an IonTorrent S5 (Thermo Fisher) machine at the Institute of Forensic Medicine, University of Zurich.

#### Analysis of the amplicon sequence data

Raw DNA sequence read data was exported to fastq files according to sequencing barcodes with the FileExporter plugin of the Torrent Suite software version 5.10. The sequence data was deposited in the NCBI Sequence Read Archive under submission number SUB6412194. Primer sequences were removed from the end of the sequences of each fastq file using the cutadapt algorithm ^58^ implemented on the Galaxy server ^59^. Trimmed sequences were quality-filtered using Usearch ^60^ with maximum expected error threshold of 100 and clustered into operational taxonomic units (OTUs) with Uparse ^61^ at 97 % minimal identity threshold and minimal OTU size of 10 sequence reads, as default settings. In some cases, these settings were slightly modified for more relaxed quality filtering and clustering to allow OTU creation for samples with lower quality sequence reads. Sequences of the resulting LR and MSH OTUs were aligned and the alignments were concatenated in Geneious version 11.1.5 (https://www.geneious.com). Our concatenated LR-MSH sequence alignment was added to the LR-MSH alignment of reference samples of the eight precious coral species listed in Table 1. The taxonomic identity of our sequences was determined by constructing a Bayesian phylogenetic tree as described in Supporting Methods S2. We noticed that several of DNA sequences obtained from the coral skeletons did not cluster in monophyletic clades with sequences of single precious coral species. We therefore performed an additional phylogenetic analysis with identical settings, which included the orthologous LR-MSH DNA sequences of all Coralliidae specimens from Tu, et al.^47^ that were identified to the species level (Supplementary Table S9).

## Supporting information

Lendvay et al. Supplementary Information

Lendvay et al. Supplementary Table S1

Lendvay et al Supplementary Table S2

Lendvay et al Supplementary Table S9

## Acknowledgements

Enzo Liverino Srl (Torre del Greco, Italy) provided some of the precious coral material for the SSEF coral collection, which we used in this study. This study benefited largely from discussions with Dr. Nozomu Iwasaki (Rissho University, Japan).

## Data availability

Raw DNA sequence data generated for this study are deposited in the NCBI Sequence Read Archive. Data used for the analyses is available as Supplementary Information.

## Supplementary Information

Supplementary Results S1. Quantitative real-time PCR results.

Supplementary Results S2. DNA sequencing results.

Supplementary Table S3. Description of the eight precious coral specious coral species relevant in the jewelry industry: distribution range and depth, taxonomy, color, trade names, harvesting history and conservation status.

Supplementary Methods S4. A – Development of taxonomic identification markers. B – Accession numbers of the DNA sequences used for taxonomic marker development. C – Majority-rule Bayesian phylogenetic tree constructed on DNA sequence data of the 29 reference Corallidae and three outgroup samples.

Supplementary Methods S5. Laboratory protocols to prevent contamination and tests performed to verify authenticity of our results.

Supplementary Table S6. Size characteristics and amount of material aliquot amounts of the 25 worked precious coral samples used both for taxonomic assignment and testing purity and quantity of five different DNA extraction methods.

Supplementary Methods S7. Real-time quantitative optimization steps, sequences of the synthetic control DNA molecules and primers used in the experiments.

Supplementary Table S8. Size characteristics of the 25 worked precious coral samples used both for DNA quantification and taxonomic assignment using “quasi non-destructive” DNA extraction technique.

Supplementary Table S9. NCBI GenBank accession numbers and sample collection data of the additional Coralliidae sequences used for the extended phylogenetic analysis.

## Author contributions

LB, AK, MSK and LEC conceived the study. JBM conducted preliminary DNA extraction and sequence analysis. LB, NVM and MG performed the laboratory work and analyzed the data. LB and LEC wrote the manuscript with support from the other co‐authors. All authors read and approved the final version of the manuscript.

