## Supplementary material for "DNA fingerprinting: an effective tool for taxonomic identification of processed precious corals": Lendvay et al. Supplementary Information

**Supplementary Results**

Supplementary Results S1.

Quantitative real-time PCR results. Available as separate Excel table.

Supplementary Results S2.

DNA sequencing results. Available as separate Excel table.

**Supplementary Methods**

Supplementary Table S3.

Description of the eight precious coral species relevant in the jewelry industry: distribution range and depth, taxonomy, color, trade names, harvesting history and conservation status.

| *Corallium japonicum* (Kishinouye, 1903) |
|---|
| Range and depth: Japan and Taiwan at the depths of 80-300 m [1,2]. |
| Taxonomy: Genetically most similar to *Corallium rubrum.* Previously assigned to the genus *Paracorallium*, but following genetic evidence [3,4] moved to the *Corallium* genus. |
| Color: Dark red with white axial core. |
| Trade names: Aka, Moro, Oxblood, Red blood. |
| Harvesting: *Corallium japonicum* fishing reached its peak in the mid-1980s (c. 10 t per year), followed by a decline in the harvest that stabilized at c. 3 t annually. In the past three decades, three quarters of *C. japonicum* was fished in Japan and the remainder in Taiwan [1]. |
| Conservation: The species is listed in Appendix III of CITES. In Japan, *Corallium japonicum* is listed as Near Threatened on the Red List of Threatened Species in Japan and the harvesting is controlled by the local prefectural governments [1]. In Taiwan, the harvest is precisely and strictly controlled by law. |

| *Corallium rubrum* (Linnaeus, 1758) |
|---|
| Range and depth: widely distributed in the western part of the Mediterranean sea [5] and also present in the Eastern Atlantic [6,7]. The species inhabits depths of a range from shallow waters of 10 m to the depth of 800 m [8], however mostly common between 20-200 m [5]. |
| Taxonomy: Genetically most similar to *Corallium japonicum*. |
| Color: Red with shades of orange, pink and "smoked" orange color. |
| Trade names: Mediterranean, Sardegna, Sardinian, Sciacca. |
| Harvesting: This species is the most prominent in the industry in terms of color, name, and quality and history [9,10]. The yearly catch of *C. rubrum* was 30-40 t in the decade of 2000-2010 and this rose to >50 t per year in the years following 2010 (FAO Fisheries Statistics Programme, http://www.fao.org), from which the majority originates from Italy, Tunisia and France [5]. |
| Conservation: Despite exploitation over millennia, *C. rubrum* is still common; however, particularly in the shallow waters, its populations are dominated by only small colonies to the extent, that this species is considered ecologically extinct [11]. For shallow populations the rising see temperatures and water acidification pose further threats under current climate change [12]. Deep-dwelling *C. rubrum* populations have remained in a more natural state than shallow living populations exhibiting higher proportion old colonies and less density [13]. *Corallium rubrum* considered as an Endangered species by the International Union for Conservation of Nature (IUCN) is listed in several international conventions: EC Habitat Directive (92/43/EEC—Appendix V); Bern Convention (Appendix III); Protocol concerning Specially Protected Areas and Biological Diversity in the Mediterranean (Barcelona Convention; Appendix III). Fishery and conservation management of the species is managed by the General Fisheries Commission for the Mediterranean [5]. |

Supplementary Table S3. continued

| *Hemicorallium laauense* (Bayer, 1956) |
| --- |
| Range and depth: Emperor Seamounts 700-2000 m [2,9] |
| Taxonomy: In the scientific and gemmological literature Deep-sea Midway coral was referred as to the scientifically undescribed coral species *Corallium sp. nov.* e.g. [9,10,14,15] and has only recently been identified as *H. laauense* [2,16], however without any documented evidence in the scientific literature. *Hemicorallium laauense* and *H. regale* are difficult to identify and were sometimes considered conspecific [17-19]. *Hemicorallium laauense* is regarded as a non-monophyletic taxon with a cosmopolitan distribution [3], which probably includes cryptic species [4]. No scientific surveys focused on *H. laauense* prior to inception of the fishery or during the period of active exploitation Science Working [20]. CIBJO suggests *Hemicorallium laauense* be traded under the name *Corallium secundum* [2,16]. |
| Color: Various colors; bright white, pink, pomegranate with red veins or spots. |
| Trade names: Deep sea, Deep-sea Midway, New coral, Sensei. |
| Harvesting: Harvesting of Deep-sea Midway coral started in 1981, and 600 t were fished between 1983 and 1986, but by the end of the decade the landings ceased due to the depletion of the fields [21]. |

| *Hemicorallium regale* (Bayer, 1956) |
| --- |
| Range and depth: Hawaii islands, 350-600 m [16,22] |
| Taxonomy: *Hemicorallium regale* and *H. laauense* are difficult to identify and sometimes were considered conspecific [17-19]. Based on their morphology, *H. regale* and *H. sulcatum* are also very closely related species [23]. CIBJO suggests that *H. regale* be traded using the name *Corallium secundum* [2,16]. |
| Color: Pomegranate-color with different intensity shades of uniform pink. |
| Trade name: Garnet. |
| Harvesting: Grigg et al. [24] reports exclusively a limited harvest of *C. regale* in Hawaii between 1999-2001, but [10] reports its fishing around the Hawaiian and Midway islands in 1970 and [16] around Hawaii in 1979, probably owing to the above mentioned similarity of these species and the confusion in the nomenclature and taxonomy of the closely related *H. laauense* (see below). |
| Taxonomy: *Hemicorallium regale* is highly similar to *H.sulcatum* [25]. |

| *Hemicorallium sulcatum* (Kishinouye, 1903) |
| --- |
| Range and depth: Japan, Taiwan and the Philippines between the depths of 100-300 m [2]. |
| Taxonomy: Miss coral has only recently been identified as the long ago described *Hemicorallium sulcatum* [23]. Based on their morphology, *H. sulcatum* and *H. regale* are very closely related [23]. CIBJO suggests *H. sulcatum* be traded using the name *Corallium secundum* [2,16]. |
| Color: uniform color, pink to violet. |
| Trade names: Miss, Missu, Misu. |
| Harvesting: The species was extensively harvested from the 1970s to the 1990s in Taiwan [23]. |

Supplementary Table S3. continued

| *Pleurocorallium elatius* (Ridley, 1882) |
| --- |
| Range and depth: Japan, Taiwan, Philippines, Indonesia, Vietnam, China at depths between 50-350 m [1,2,22]. |
| Taxonomy: Very closely related to *Pleurocorallium konojoi*; no genetic difference discovered in the DNA regions studied by Tu, et al. [4], and only nine bp difference discovered between the entire mitochondrial DNA genome of the two species differs by 10 bp [26]. |
| Color: Flesh pink with different color intensities from bright red, salmon, orange to flesh, usually with white axial core. |
| Trade names: Angel skin, Boké, Cerasuolo, Magai, Momo, Pelle d'angelo, Satsuma. |
| Harvesting: *Pleurocorallium elatius* has been the most harvested species of the Western Pacific; its harvest peaked in 2012 c. 26.5 t, [1]. Recently, *P. elatius* was fished mainly in waters of China (12-20 t per year), followed by Taiwan and Japan (c. 5.5 t per year both). |
| Conservation: The species is listed in Appendix III of CITES. In Japan, *Pleurocorallium elatius* is listed as Near Threatened on the Red List of Threatened Species in Japan and the harvesting is controlled by the local prefectural governments [1]. In Taiwan, the harvest is precisely and strictly controlled by law. In China, it is protected by law, and its fishing is legally not allowed. |

| *Pleurocorallium konojoi* (Kishinouye, 1903) |
| --- |
| Range and depth: Japan, Taiwan, China, Vietnam, Philippines, 50-380 m [1,2,9]. |
| Taxonomy: Very closely related to *Pleurocorallium elatius*; no genetic difference discovered in the DNA regions studied by Tu, et al. [4], and only 10 bp difference discovered between the entire mitochondrial DNA genome of the two species [26]. |
| Color: Milky white and red or pink speckled white. |
| Trade names: Bianco, Shiro, Pure white. |
| Harvesting: The global annual harvest of *Pleurocorallium konojoi* is less than 1 t. The main producers are Japan and Taiwan with over 90% of the catch [1]. |
| Conservation: The species is listed in Appendix III of CITES. In Japan, *Pleurocorallium konojoi* is listed as Near Threatened on the Red List of Threatened Species and the harvesting is controlled by the local prefectural governments [1]. In Taiwan, the harvest is precisely and strictly controlled by law. In China, it is protected by law, and its fishing is legally not allowed. |

| *Pleurocorallium secundum* (Dana, 1846) |
| --- |
| Range and depth: Emperor seamount, Hawaiian Archipelago, 340-600 m [2,9] |
| Color: Red speckled or veined white or pink; uniform clear pink. |
| Trade names: Midway, Rosato, White/Pink |
| Harvesting: The commercial harvesting of the species began in 1965 and nearly 230 t were fished until 1970 [24,27]. The species has not been harvested since 1990, except for minor catches in 2015 and 2016 [1]. |
| Conservation: The species is listed in Appendix III of CITES. |

Supplementary Methods S4.

A – Development of taxonomic identification markers.

We developed two markers with the aim to be able to distinguish the eight precious coral species listed by CIBJO as relevant for the jewelry industry. Sequence alignments of the eight precious coral species were created for the six mitochondrial DNA fragments analyzed by Tu, et al. [4]. The alignments contained sequences of altogether 29 specimens with each species represented by at least two sequences (see Supplementary Methods S4B). Sequence alignments were performed in Geneious version 11.1.5 with default settings. We searched for maximum 200 bp long variable regions in the alignments that were flanked by conserved regions that allow primer development without mismatches on the priming sites.

Following visual inspection of the alignments, and testing the phylogenetic resolution of multiple target regions, we selected highly variable regions of the large ribosomal RNA gene subunit (LR) and the putative mismatch repair protein (MSH), respectively. Flanking both selected gene fragments, primer-pairs were designed with Primer3 version 2.3.7. [28] implemented in Geneious. The primer design was set to flank ca. 150 bp long regions on sites uniform in all aligned sequences and resulted in the primers LR-F 5'TTCATCACAGTGAGGGTTTGT3' and LR-R 5'TGCAAAGAAGGAGAACAAAAGG3' for the LR gene and MSH-F 5'CGAAAGCGGATAAAAGCTACC3' and MSH-R 5'CCTCACTGTCAGGCTAATGAG3' for the MSH gene, respectively.

Both the LR and MSH regions flanked by the designed primers were aligned to the *Corallium rubrum* mitochondrial genome (GenBank accession number AB700136) as reference. The LR fragment stretches from the 140$^{th}$ to the 293$^{rd}$ position of the complete LR gene and is 154 bp long in *Corallium* and *Hemicorallium*, and 128-129 bp long in *Pleurocorallium*, respectively. The MSH gene fragment stretches from the 706$^{th}$ to the 841$^{st}$ position of the complete MSH gene and is uniformly 136 bp long in all aligned 29 precious coral sequences.

Orthologous sequences of three *Paragorgia sp.* specimens used by Tu, et al. [4] were then included in the alignments as outgroup, and taxonomic resolution and phylogenetic tree topology was tested based on the LR and MSH sequences (see Supplementary Methods S4B). Incongruence length difference test as partition homogeneity test with heuristic search and parsimony was run in PAUP version 4.0 [29] [30]. No incongruence was found between the LR and MSH datasets ($p = 0.81$), therefore the two regions were concatenated and Bayesian phylogenetic tree were created with partitioning the two genes using MrBayes version 3.2.7 [31].

Following the recommendation of the MrBayes software manual, instead of applying an a priori set substitution model, we set the "mixed substitution model prior", letting the software to sample across nucleotide substitution models. The analysis was run $10^7$ generations sampling every $10^3$ generations and 25 % burn-in. Majority rule consensus tree was visualized in FigTree [32].

The resulting tree reveals the three main phylogenetic clades of the Corallidae family, i.e. Clade Ia – *Corallium*, Clade Ib – *Hemicorallium* and Clade II – *Pleurocorallium* (Supplementary Results S4C, see [3,4]). Specimens of each species cluster together in a monophyletic group with two exceptions. The two samples of *Pleurocorallium laauense* – although identical to each other and different from all other sequences – form an unresolved clade basal to the other *Hemicorallium* species. The samples of *Pleurocorallium elatius* and *P. konojoi*, which have identical LR-MSH sequences, form an unresolved tree branch. Distinguishing the latter two species was impossible based on any other subsets of the data set of Tu, et al. [4].

To demonstrate the ability of our two short markers to reconstruct the phylogenetic relationships among the eight precious coral species, we built a phylogenetic tree using sequence data of all six mitochondrial regions of the 32 reference samples (see Supplementary Methods S4B). Phylogenetic analysis on the 3620 bp long sequence alignment was performed in the same way as described above.

B – Accession numbers of the DNA sequences used for taxonomic marker development.

| Taxon | 16S | ND1 | ND2-16S | ND6-ND3 | MSH | IGR1 |
|---|---|---|---|---|---|---|
| *Corallium japonicum*-1 | AB595189 | AB595189 | AB595189 | AB595189 | AB595189 | AB595189 |
| *Corallium japonicum*-2 | KF850198 | KF854776 | KF850283 | KF850341 | KF854882 | KF855041 |
| *Corallium japonicum*-3 | KF850193 | KF854785 | KF850285 | KF850345 | KF854891 | KF855044 |
| *Corallium japonicum*-4 | JX647961 | KF854787 | JX648084 | JX840706 | KF854892 | KF855045 |
| *Corallium japonicum*-5 | KF850190 | KF854788 | KF850286 | KF850347 | KF854894 | KF855046 |
| *Corallium rubrum*-1 | AB700136 | AB700136 | AB700136 | AB700136 | AB700136 | AB700136 |
| *Corallium rubrum*-2 | KF286554 | KF854765 | KF286555 | KF286556 | KF854871 | KF855039 |
| *Hemicorallium laauense*-1 | JX647898 | KF854832 | JX648023 | JX840647 | KF854938 | KF855002 |
| *Hemicorallium laauense*-2 | KF850229 | KF854761 | KF850271 | KF850333 | KF854867 | KF855026 |
| *Hemicorallium regale*-1 | JX647901 | KF854826 | JX648026 | JX840650 | KF854932 | KF855000 |
| *Hemicorallium regale*-2 | JX647893 | KF854789 | JX648018 | JX840642 | KF854895 | KF854987 |
| *Hemicorallium regale*-3 | JX647883 | KF854736 | JX648008 | JX840632 | KF854842 | KF854984 |
| *Hemicorallium regale*-4 | JX647892 | KF854823 | JX648017 | JX840641 | KF854929 | KF854999 |
| *Hemicorallium regale*-5 | KF850196 | KF854778 | KF850284 | KF850343 | KF854884 | KF854977 |
| *Hemicorallium sulcatum*-1 | KF850191 | KF854795 | KF850269 | KF850349 | KF854901 | KF854988 |
| *Hemicorallium sulcatum*-2 | KF850187 | KF854794 | KF850296 | KF850348 | KF854900 | KF854990 |
| *Pleurocorallium elatius*-1 | KF850201 | KF854772 | KF850258 | KF850301 | KF854878 | KF855028 |
| *Pleurocorallium elatius*-2 | KF850189 | KF854790 | KF850245 | KF850304 | KF854896 | KF855036 |
| *Pleurocorallium elatius*-3 | KF850188 | KF854791 | KF850263 | KF850305 | KF854897 | KF855030 |
| *Pleurocorallium konojoi*-1 | AB595190 | AB595190 | AB595190 | AB595190 | AB595190 | AB595190 |
| *Pleurocorallium konojoi*-2 | KF850202 | KF854771 | KF850257 | KF850300 | KF854877 | KF855034 |
| *Pleurocorallium konojoi*-3 | KF850203 | KF854770 | KF850256 | KF850299 | KF854876 | KF855033 |
| *Pleurocorallium konojoi*-4 | KF850200 | KF854774 | KF850259 | KF850302 | KF854880 | KF855029 |
| *Pleurocorallium konojoi*-5 | KF850199 | KF854775 | KF850260 | KF850303 | KF854881 | KF855035 |
| *Pleurocorallium konojoi*-6 | KF850186 | KF854796 | KF850264 | KF850306 | KF854902 | KF855031 |
| *Pleurocorallium konojoi*-7 | KF850184 | KF854800 | KF850266 | KF850307 | KF854906 | KF855032 |
| *Pleurocorallium secundum*-1 | JX647953 | KF854817 | JX648076 | JX840698 | KF854923 | KF854967 |
| *Pleurocorallium secundum*-2 | JX647952 | KF854757 | JX648075 | JX840697 | KF854863 | KF854956 |
| *Pleurocorallium secundum*-3 | KF850194 | KF854784 | KF850238 | KF850310 | KF854890 | KF854951 |
| Outgroup samples: | | | | | | |
| *Paragorgia* sp-1 | KC782351 | KC782351 | KC782351 | KC782351 | KC782351 | KC782351 |
| *Paragorgia* sp-2 | KC782350 | KC782350 | KC782350 | KC782350 | KC782350 | KC782350 |
| *Paragorgia* sp-3 | KC782349 | KC782349 | KC782349 | KC782349 | KC782349 | KC782349 |

C – Majority-rule Bayesian phylogenetic tree constructed on DNA sequence data of the 29 reference Corallidae and three outgroup samples. On the top, tree constructed from the 293 bp long sequence alignment of the combined LR-MSH regions. On the bottom, tree constructed from 3620 bp long sequence alignment of six mitochondrial genes of the same samples. Note the similarity of the tree topologies.

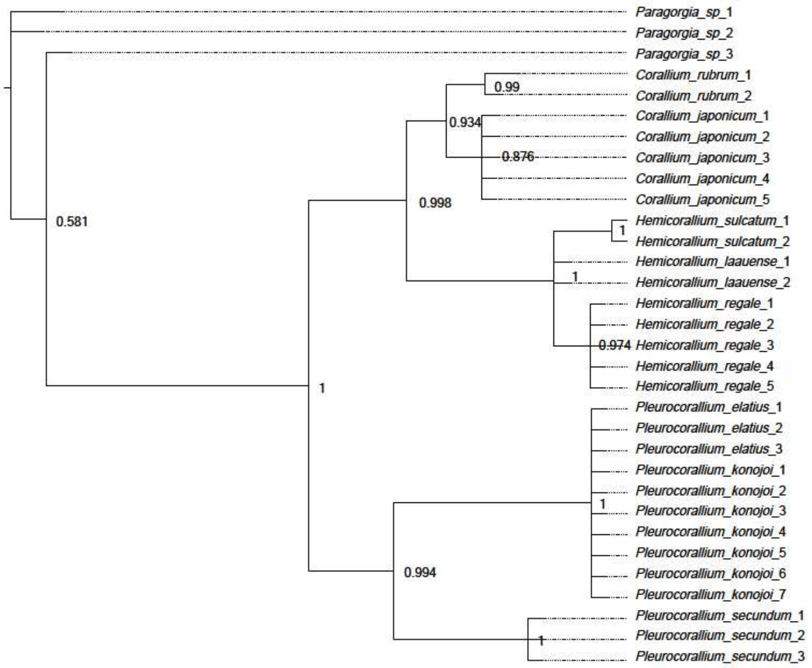


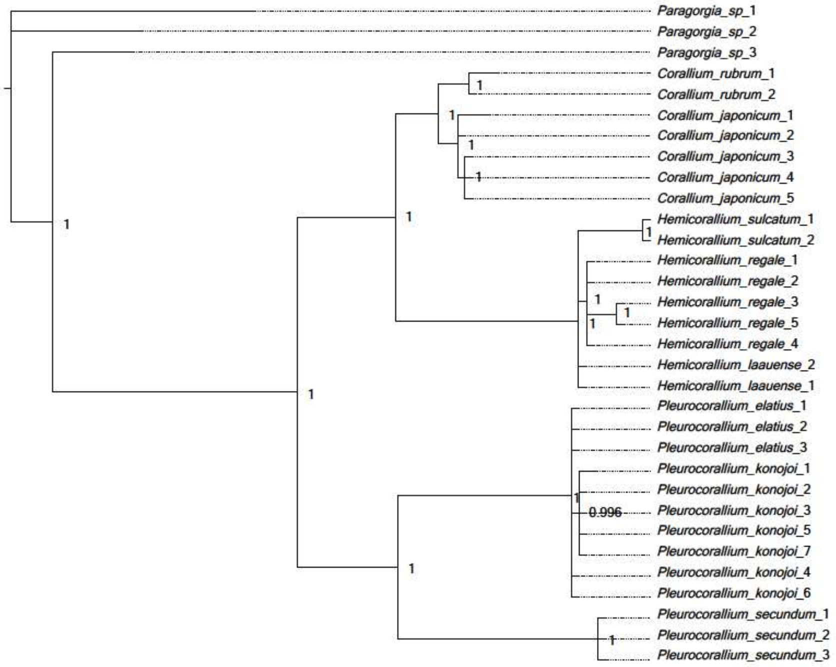

Supplementary Methods S5.

Laboratory protocols to prevent contamination and tests performed to verify authenticity of our results.

We applied rigorous measures to prevent introduction of laboratory contamination. Prior to being processed, coral skeletal axis samples were thoroughly cleaned with 5% neodisher lm3 (Dr. Wiegert, Hamburg, Germany), an alkaline surfactant solution. We refrained from applying the standard method of cleaning sample surfaces by bleaching, as this might deteriorate the coloration of our sensitive samples. Pulverization of the coral skeletons was performed in a laboratory room separated from the room where the remaining steps of the DNA extraction was carried out. We took particular care to avoid sample cross-contamination during the pulverization step and the re-use of the mortars. After each use, mortars, pistils and scalpels used for pulverization and sample collection were first immersed in 2% hydrochloric acid for 10 minutes to dissolve remaining uncollected coral powder. Then, these utensils were thoroughly washed with soap and neodisher lm3 solution, followed by rinsing with deionized water and 70% ethanol. The drill heads were cleaned after each sample in the same way.

To provide evidence that this cleaning procedure was effective enough to prevent carryover of coral material that would cause detectable DNA contamination in the following sample, we pulverized material of sterile plastic lab consumable pieces identically to the method used for the corals. Then, the DNA extraction protocol was implemented using the plastic samples identically as for as the coral samples. Altogether eight plastic control sample extractions were performed.

To prevent the dispersal of airborne coral powder during the sample pulverization, the mortars were covered with a rubber glove during the crushing of the samples. Furthermore, to prove that airborne coral powder did not cause detectable DNA contamination, we placed open-lid plastic tubes filled with water in the proximity of the sample being crushed or drilled. The water sample was then processed through the subsequent DNA extraction process. Altogether eight open-lid water control extractions were performed.

For each set of maximal 13 DNA samples, two additional blank extraction control samples were processed in the extraction protocol starting from the lysis step.

Ingredients of the qPCR reactions were assembled in a pipetting hood in a pre-PCR room. After setting up a PCR, the interior of the hood and pipettes were cleaned with neodisher lm3 and the hood was irradiated with UV light for at least 30 minutes. Three non-template controls were included on each 96-well reaction plate. For the absolute DNA quantification reactions, these non-template controls were placed adjacent to the wells of the most concentrated standard reaction samples ($10^7$ DNA copies) to prove that cross-contamination between the wells was not present. The PowerUP SYBR Green Master-mix used for amplification incorporates uracils, and has a built-in enzyme to degrade uracil-containing DNA at the beginning of the PCR (detailed in Supplementary Methods S7).

Throughout our experiments, none of the extraction controls (pulverized plastic, open-lid water or blank controls) or PCR non-template controls amplified to the extent to be detected by the qPCR analysis software.

Melting-curves of the triplicate qPCR reactions were compared by eye to check that the identical DNA molecule has been amplified in each of them. For the standard molecules in the qPCR reactions (quantification standard in the absolute quantification assays and the internal amplification control in the inhibition-tests), we checked that the melting-curve matches with the theoretical melting-curve projected for the given standard molecule DNA sequence by the uMelt software [33].

For both the LR and MSH marker, the DNA sequence pertaining the OTU with the largest sequence read number was subjected to NCBI megaBLAST search. We verified that all sequences match sequences of the Coralliidae family, and that there is no phylogenetic contradiction between the hits with highest similarity obtained for the LR and MSH markers, respectively.

Supplementary Table S6.

Size characteristics and amount of material aliquot amounts of the 25 worked precious coral samples used both for taxonomic assignment and testing purity and quantity of five different DNA extraction methods.

| No | | Weight (mg) | Longest side (mm) | Material aliquot (mg) |
|---|---|---|---|---|
| 1 | 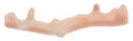 | 1.15 | 35 | 100 |
| 2 | 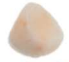 | 0.80 | 9 | 100 |
| 3 | 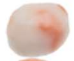 | 1.70 | 12 | 100 |
| 4 | 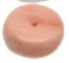 | 1.64 | 12 | 100 |
| 5 | 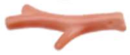 | 0.45 | 22 | 75 |
| 6 | 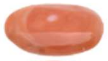 | 0.95 | 17 | 100 |
| 7 | 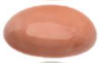 | 2.24 | 19 | 100 |
| 8 | 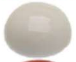 | 2.59 | 15 | 100 |
| 9 | 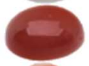 | 0.96 | 12 | 100 |
| 10 | 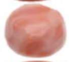 | 1.86 | 12.5 | 100 |
| 11 | 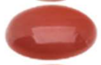 | 0.56 | 12 | 95 |
| 12 | 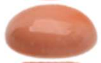 | 1.82 | 17 | 100 |
| 13 | 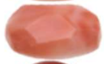 | 1.06 | 15 | 100 |
| 14 | 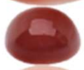 | 1.07 | 12 | 100 |
| 15 | 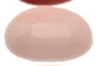 | 1.31 | 14.5 | 100 |
| 16 | 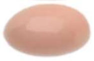 | 0.96 | 14 | 100 |
| 17 | 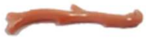 | 0.82 | 12 | 100 |
| 18 | 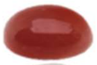 | 0.99 | 14 | 100 |
| 19 | 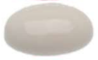 | 0.74 | 10 | 100 |
| 20 | 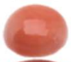 | 0.57 | 10 | 90 |
| 21 | 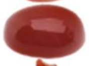 | 0.99 | 11.5 | 100 |
| 22 | 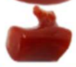 | 0.85 | 18 | 100 |
| 23 | 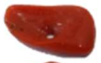 | 1.00 | 14 | 87 |
| 24 | 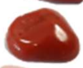 | 0.64 | 36 | 100 |
| 25 | 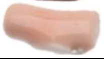 | 0.71 | 13 | 100 |

Supplementary Methods S7.

Real-time quantitative optimization steps, sequences of the synthetic control DNA molecules and primers used in the experiments.

Optimal DNA template input amount was selected based on a subset of the coral DNA extracts representing different extraction methods. Input DNA extract volumes of 1 µl, 3 µl and 5 µl were tested, and the success of amplification and the shape of amplification curves were observed. The selected 3 µl DNA extract generally produced amplicons, and PCR amplification curves appeared with sigmoidal shape indicating no signs of "flattening" of the curves due to PCR inhibition. Identical amount (3 µl) of coral DNA extract was then used as input in the assays testing the PCR inhibition effect of the coral DNA extracts.

The template copy number of the standard quantification molecule used in the inhibition tests was selected based on preliminary coral DNA absolute quantification results. Our aim was to set the template amount to the order of magnitude present in the coral DNA extracts to make the inhibition tests realistic. We selected $10^3$ molecules template copy number as template number of the standard quantification molecule, because we considered this amount high enough to produce robust amplification, whereas it does not exceed the amount of template molecules found in 3 µl of coral skeleton DNA extracts.

Optimal primer annealing temperatures of the newly developed primers were selected based on agarose gel electrophoresis of gradient PCR products. The chosen annealing temperature, 56 °C, produced strong amplification of the target fragment without unspecific of primer-dimer amplification.

Primer concentrations were chosen after testing 25 combinations of forward-reverse primer concentrations covering the range of the potential primer concentrations suggested by the user manual of the PowerUp SYBR Green Master Mix. In particular, qPCR reactions were run with different combinations of forward and reverse primer concentrations ranging from 5 µM to 15 µM, respectively, and the primer concentrations were chosen where the amplification of the same template amplified with the lowest Ct value. 15 µM concentration of both forward and reverse primers producing the earliest amplification.

At the beginning of each amplification, Uracil N-glycosylase (UNG) activation step was applied for two minutes at 50 °C following the vendor's recommendations, to activate the built-in UNG enzyme of the PowerUp SYBR Green Master Mix. UNG cleaves DNA molecules containing uracil molecules incorporated by the master-mix, therefore reducing the possibility of carryover contamination from previous qPCR runs.

We tested the dynamic range of the calibration curve used for the absolute quantification of the coral DNA extracts. Standard reactions were prepared with GBlocks synthetic oligonucleotides used as template DNA with copy numbers ranging from $10^1$ to $10^8$ with 10-fold dilutions in triplicates. Each dilution step performed by adding 45 µl water to 5 µl of oligonucleotide solution. We inspected whether amplifications were any unamplified reactions and the coefficient of determination ($R^2$ value) of the calibration curve. We observed that that the reactions reliably amplified throughout the tested range of template input and $R^2$ value was always >0.99. We therefore considered the entire range of $10^1$ to $10^8$ template input within the dynamic range of the qPCR assay.

The quantitative real-time PCR reactions were analyzed with automatic baseline threshold settings in the HD Real-time PCR analysis software version 1.2 (Thermo Fisher).

Sequence of the internal amplification control molecule:
5'AACTTGGCTTTAATGGACCTCCAGGGATTAATAGGGTGCCACAATAGTGGCGCA CCTTTAGGTGCCATGTGCGCGCATAGCCCGTGGCACCCTATGGAATTAACACTAG GGCTTATTATACTAATACTGTTAACGTATGGATTAAAGGGCGCGACTCTAAGATT AGCAATATTCTGGTGCCATGTAAGGATGAATGT3'

Forward primer "SPUD-F":
5'AACTTGGCTTTAATGGACCTCCA3' [34]

Reverse primer "SPUD-R":
5'ACATTCATCCTTACATGGCACCA3' [34]

Sequence of the quantification standard molecule:
5'TTCATCACAGTGAGGGTTTGTCTTATAACGATTAGAATCTCAGCCCGGGAAATT AGGTATGCAATTGTGATAATCATATTATTCGGGATCACAATTAAGGAATCCCACG GTCCCCGATACGCGGGTGTACCGTGGATTTCCACGCGGTGAAAACACCCTGGCAT AATTAGGGTTTGGAAAACAAGAGGAAGAAACGT3'

Supplementary Table S8.

Size characteristics of the 25 worked precious coral samples used both for DNA quantification and taxonomic assignment using “quasi non-destructive” DNA extraction technique.

| No | | Weight (mg) | Longest side (mm) |
|---|---|---|---|
| 26 | 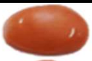 | 0.33 | 10 |
| 27 | 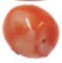 | 1.22 | 9 |
| 28 | 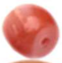 | 0.15 | 4 |
| 29 | 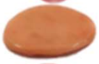 | 1.08 | 18 |
| 30 | 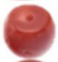 | 0.3 | 6 |
| 31 | 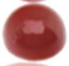 | 0.43 | 9 |
| 32 | 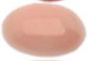 | 0.38 | 10 |
| 33 | 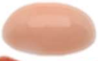 | 1.55 | 17 |
| 34 | 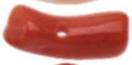 | 0.1 | 8 |
| 35 | 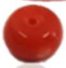 | 0.21 | 6 |
| 36 | 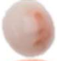 | 0.49 | 8 |
| 37 | 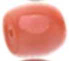 | 0.27 | 6 |
| 38 | 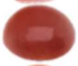 | 0.37 | 9 |
| 39 | 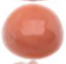 | 0.44 | 8 |
| 40 | 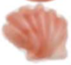 | 0.38 | 10 |
| 41 | 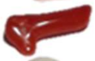 | 0.12 | 10 |
| 42 | 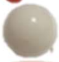 | 0.45 | 8 |
| 43 | 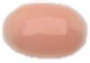 | 0.31 | 9 |
| 44 | 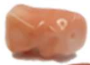 | 1.02 | 13 |
| 45 | 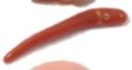 | 0.14 | 19 |
| 46 | 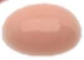 | 0.34 | 9 |
| 47 | 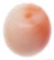 | 0.56 | 7 |
| 48 | 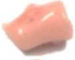 | 0.57 | 10 |
| 49 | 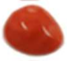 | 0.68 | 13 |
| 50 | 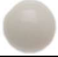 | 0.48 | 8 |

Supplementary Table S9.

NCBI GenBank accession numbers and sample collection data of the additional Coralliidae sequences used for the extended phylogenetic analysis. Data extracted from the publication of Tu et al. [4]. Available as separate Excel table.
